# Brain plasticity for visual words: Elementary school teachers can drive changes in weeks that rival those formed over years

**DOI:** 10.1101/2024.09.17.613570

**Authors:** Fang Wang, Elizabeth Y. Toomarian, Radhika S. Gosavi, Blair Kaneshiro, Anthony M. Norcia, Bruce D. McCandliss

## Abstract

This study aims to investigate the impact of vocabulary acquisition through short-term classroom learning and its relation to broader forms of vocabulary learning through long-term exposure in daily life. Through a two week of “learning sprint” in collaboration with a local elementary school and EEG-Steady State Visual Evoked Potentials (EEG-SSVEP) paradigm, we assessed new vocabulary learning in first and second graders within their pedagogical environment. We then compared the results with the word frequency effect, a well-established phenonmenon that reflects long-term vocabulary learning. After two weeks of classroom instruction, newly acquired words elicited neural responses similar to those of high-frequency words, with the effect significantly correlated with children’s phonological decoding skills. Additionally, we successfully replicated the word frequency effect using the SSVEP paradigm for the first time. These findings highlight the potential of the “learning sprint” model for conducting neuroscience research in authentic educational settings, thereby fostering a stronger connection between education and neuroscience.

## 1 Introduction

Vocabulary plays a pivotal role in both communication and academic success. Hence, it is important to understand the neuromechanisms underlying vocabulary learning especially in early readers. Children’s vocabularies grow by thousands of words each year, both through incidental, accumulative encounters with words in books and verbal contexts, as well as through explicit, systematic teaching in school, where novel words’ pronunciation and meaning are directly taught to children (Robbins & Ehri, 1994).

One common method for estimating readers’ cumulative experience with words gained over their reading lifespan is word frequency. Estimates of word frequency are typically based on the number of occurrences within a large sample of text (corpus or textbook). High-frequency words are processed more efficiently than low-frequency words (Balota et al., 2004; White, 2008), a phenomenon well known as the word frequency effect (for a review, see Brysbaert et al. (2018)). Our recent work has shown that the brain signals of early readers (i.e., kindergartners to second graders) in response to visual word forms are significantly influenced by their cumulative prior experience with the specific visual words. Specifically, neural circuits exhibit greater tuning to familiar, high-frequency words compared to unfamiliar pseudowords (Wang et al., 2022).

Educational neuroscience, as an emerging sub-discipline of cognitive neuroscience, aims to bridge the gap between educational practices and neuroscience research, uncovering experience dependent brain plasticity in learning. It is thus crucial for educational neuroscience to understand not only how neural responses reflect the cumulative exposure to visual words, but also how deliberate, week-to-week, explicit efforts to teach new visual words in school drive changes in neural responses to these new words. Research in this direction can further inform teaching methods, curriculum development, and educational policies.

However, most previous research on the neuromechanisms of new vocabulary learning has primarily relied on laboratory-based efforts, typically involving multiple sessions of training in a controlled lab setting (Alharbi, 2019; Davis et al., 2009; Nation et al., 2007). To provide more relevant insights into how educational practices can be optimized to improve learning outcomes, class-based naturalistic teaching practice in school offers greater relevance than computer-based learning in a laboratory.

Thus, the main goal of the present study was to investigate how direct, short-term classroom learning drives changes in the neurocognitive processes of new vocabulary words. Specifically, we aimed to determine whether newly learned words elicit similar/different neural responses compared to words learned over a much longer period of time. Additionally, we sought to explore how the neural correlates of the word frequency effects can be reliably captured within a Steady State Evoked Potentials (SSVEP) oddball paradigm (for a review, see Norcia et al. (2015)). The high signal-to-noise ratio advantages of SSVEP facilitate rapid measurements of neural signals in just a few minutes of recording (Wang et al., 2021). This advancement over traditional Event-related Potentials (ERP) measures enables the capture of words learning effect within the school context concurrently with investigation of word frequency in the same subjects during a single session.

The entire study, including EEG data collection, was carried out within an elementary school setting. We utilized a Research-Practice Partnership between a university research initiative and a local elementary school (Gosavi & Toomarian, 2024); researchers and practitioners co-designed the study. Learning strategies and activities reflected teachers’ authentic daily practices. During the two weeks of “learning sprint”, teachers led their class in learning a list of uncommon vocabulary words 15 minutes per school day. Moreover, students were not only participants, but also played an active role in the research process through constant interactions with researchers and teachers. Cognitive processes underlying new word learning were assessed using EEG, which was recorded in a dedicated laboratory set up inside of the school (Toomarian et al., 2024). The study’s design, which randomized the selection of words for teaching while reserving others as controls, is positioned to investigate a causal link between specific learning activities and changes in word representations.

Of note, we also included high- and medium-frequency words from a corpus in the same experiment. This allows us to: (1) examine the extent to which the new vocabularies were learned in a short period of time, by comparing brain responses to newly learned words with those to high- and medium-frequency words; and (2) capture the well-established word frequency effect with the SSVEP paradigm for the first time.

This naturalistic classroom learning study provides a unique extension of existing knowledge on children’s vocabulary learning. Our goal was to reveal the causal relationships between word learning and word representations’ retrieval, as well as to replicate and extend the word frequency effect. Moreover, the study showcases the RPP paradigm together with the “learning sprint” model as a promising direction for future research in the field of educational neuroscience. Theoretically, such studies may shed new light on theories of word reading development and models of visual word recognition. On a practice level, via collaborations between researchers and educational professionals, such studies could yield insights for educational interventions and activities that promote efficient new vocabulary acquisition and improve reading fluency.

## 2 Methods

### 2.1 Participants

Forty-eight English-speaking children from three classes at an independent school participated in this two-week learning study. All participants participated a behavioral lexical decision task both before and after the learning sprint. Of these 48 participants, 30 children (ten from each class), with normal or corrected-to-normal vision and no reading disabilities, completed and EEG sessions after the classroom learning sprint. Two participants (from two classes) were excluded due to data quality issues, resulting in *N* = 28 participants ranging in age from 6.75 to 8.89 years old (*m* = 7.69 years, *s* = 0.57 years, 14 males). The 28 participants whose EEG data were analyzed included 16 first graders (*m* = 7.32 years, *s* = 0.38 years, 9 males) and 12 second graders (*m* = 8.19 years, *s* = 0.38 years, 5 males)^1^.

### 2.2 General Cognitive Assessments

Each participant participated in a 30-minute behavioral session, on average two days (s = 3 days) after the EEG session. All children were tested on handedness (Edinburgh Handedness Inventory, Oldfield (1971)), phonological awareness and rapid naming abilities using two sub-tests (Rapid Automatized Naming of letters and colors) of the Comprehensive Test of Phonological Processing, Second Edition (CTOPP-II, Wagner et al. (2013)), word reading efficiency (Test of Word Reading Efficiency, Second Edition, TOWRE-2, Torgesen et al. (2012)), and word decoding ability (Woodcock-Johnson Tests of Achievement, Fourth Edition, WJ-IV, Schrank et al. (2014). Results of the behavioral assessments are summarized in Table 1.

**Table 1:**
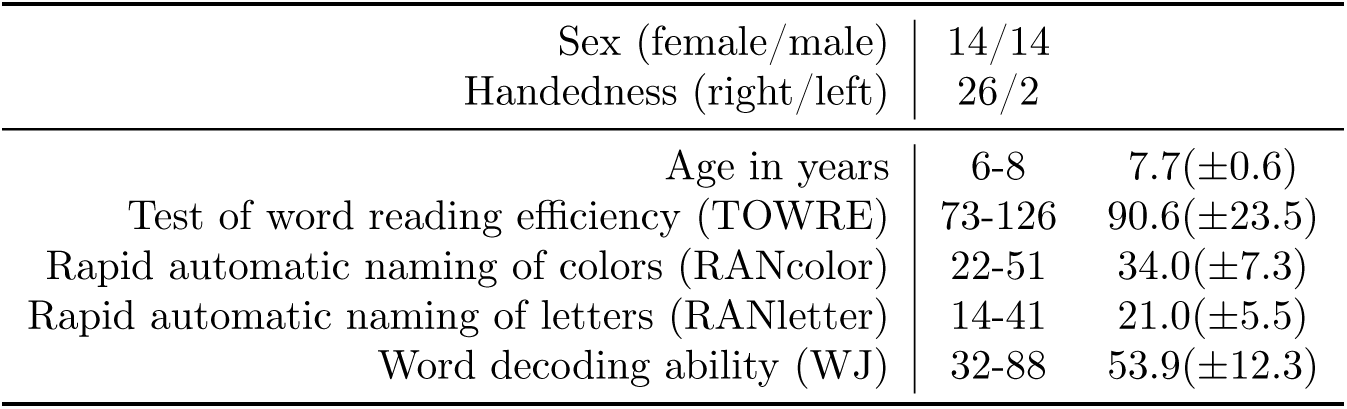
Performance on behavioral assessments. Values are range and mean(*±SD*). TOWRE: Number of real words and pronounceable nonwords read in 45 seconds. RAN: Time (ms) used to quickly and accurately name all stimuli (e.g., letters or colors) on a test form. WJ: Number of correctly named letters and words.

|  |  |  |
| --- | --- | --- |
| Sex (female/male) | 14/14 |  |
| Handedness (right/left) | 26/2 |  |
| Age in years | 6-8 | 7.7(±0.6) |
| Test of word reading efficiency (TOWRE) | 73-126 | 90.6(±23.5) |
| Rapid automatic naming of colors (RANcolor) | 22-51 | 34.0(±7.3) |
| Rapid automatic naming of letters (RANletter) | 14-41 | 21.0(±5.5) |
| Word decoding ability (WJ) | 32-88 | 53.9(±12.3) |

### 2.3 Study Procedure and Learning Sprint

Before initiating the classroom learning sprint, teachers were briefed on the concept. They engaged in collaborative brainstorming sessions to pool ideas and insights. Drawing from their collective feedback and existing teaching approaches, a tailored set of teaching strategies focusing on phonemic awareness, phonics, as well as handwriting, and a list of “magic” words were developed for the sprint. Following this, a research plan was crafted to guide the study’s launch and initiate the learning sprint (Figure 1A).

**Figure 1:**
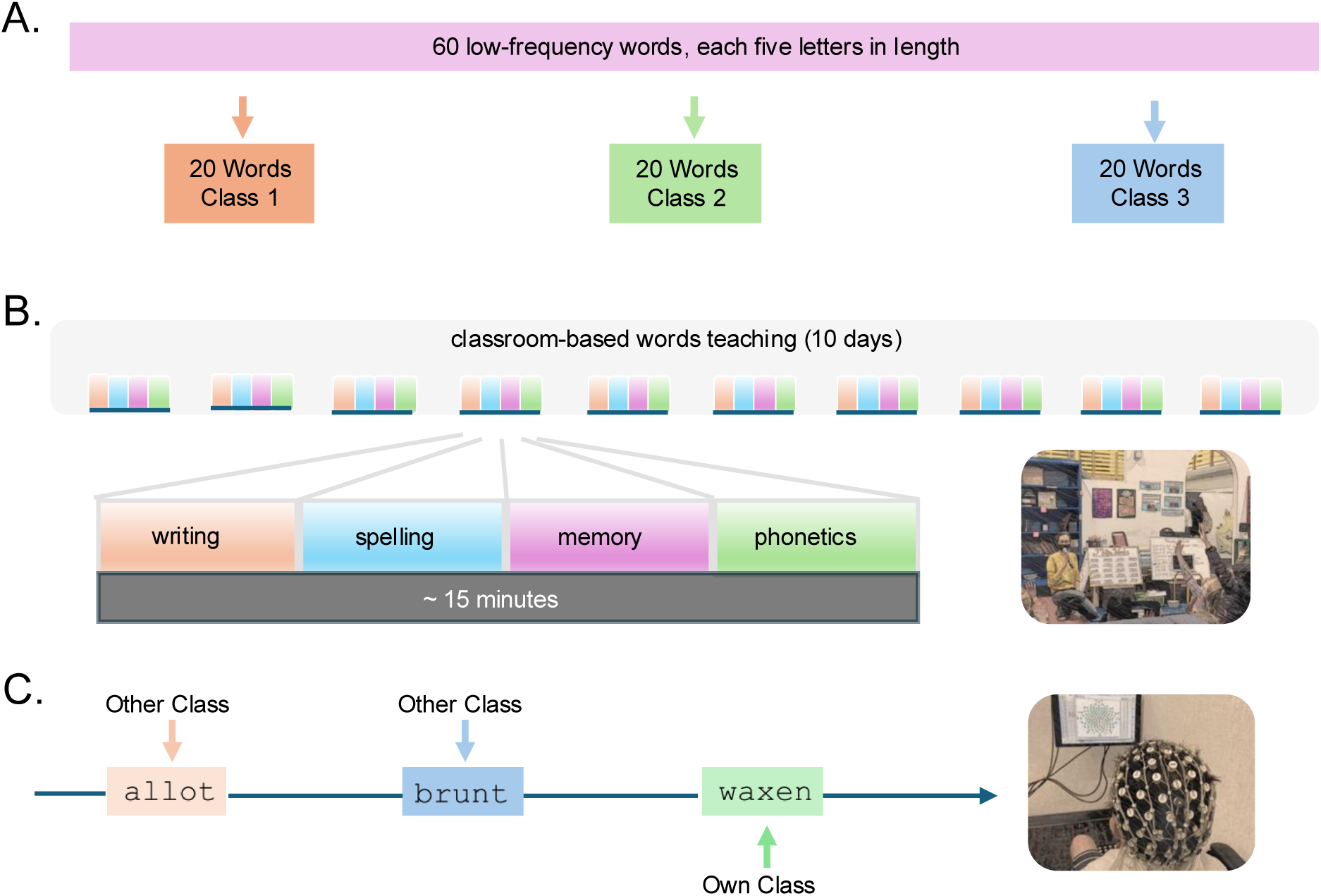
Study Procedure. A: Counterbalancing new vocabulary words. Researchers and teachers collaborated to design teaching strategies and create vocabulary lists. Sixty new low-frequency words were semi-randomly assigned to three classes (20 words per class); B: Learning sprint. During a two-week learning sprint (15 minutes daily, 5 days per week), teachers focused on phonemic awareness, phonics, writing, and memory. C: EEG recording at school. After completing their learning activities, participants individually visited the Brainwave Learning Center studio at their school for EEG recording. During the session, participants were shown words they had learned in their own class as well as words they had not learned from the other two classes.

During the two-week classroom-based learning sprint, each teacher led their class in learning a list of 20 assigned new low-frequency vocabulary words (see Section 2.4, Stimuli). Teachers collaboratively designed and implemented activities based on their classroom experiences. Display boards, featuring laminated cards labeled with the selected words and attached with velcro, served as interactive tools for teachers to dynamically engage with their students. A comprehensive tracker was created for each classroom, providing both teachers and students with a tangible visual representation of their progress over the course of the learning sprint. Participants learned the assigned word list 15–20 minutes per day, 5 school days per week, for two weeks total. The range of activities employed in the classrooms was diverse and focused on phonemic awareness, phonics and writing, encompassing vocabulary flashcards, spelling games, memory exercises, poetic exploration, Kahoot quizzes (an online game-based learning platform), and more. After each class finished their respective learning activities, participants individually visited the EEG recording studio in their school. Experimental sessions took place during the school day, allowing students to participate in research and return to class (Figure 1B).

### 2.4 Stimuli

Sixty low-frequency words (LFW, <1 per million), each five letters in length, were chosen from the MRC psycholinguistic database (Coltheart, 1981) and were semi-randomly divided into three lists, each consisting of 20 items. Unigram, bigram, trigram frequencies, number of phonemes and syllables, and orthographic neighborhood size were well matched (all *F* (2, 59) < 0.57, all *p* > 0.57) across these three word lists, which were randomly assigned across the three classes.

To better evaluate the degree of classroom-based word learning and compare short-term and long-term word learning, the study also involved high-frequency words (HFW) and medium-frequency words (MFW), all comprising five letters. The HFW (mean = 1006 per million, range 523–1821 per million) and MFW (mean = 221 per million, range 200–246 per million) were chosen from *The Educator’s Word Frequency Guide* (Zeno et al., 1995). Finally, five-letter pseudowords (PW) were included for the examination of lexical level representations compared to real words. PW were built on an item-by-item basis by shuffling letters across the set of high- and medium-frequency words used in the current study. PW were thus pronounceable and well-matched for orthographic properties (at both letter and structure level) of high- and medium-frequency words. Unigram, bigram, and trigram frequencies (all *t* < 1.98, all *p* > 0.05) and orthographic neighborhood sizes (*t*(69) = 1.29, *p* = 0.20) were matched between words and pseudowords.

In all, the stimuli set comprised 20 HFW, 20 MFW, 60 LFW, and 80 PW, for 180 exemplars total. We investigated three experimental conditions as summarized in Figure 2. In order to examine the effect of new word learning, in condition 1 (Figure 2A), learned low-frequency word deviants from a student’s own class were embedded in a stream of unlearned low-frequency words from the other two classes (LFW_L_-LFW_UL_). To examine the extent to which the new vocabularies were learned in a short period of time, two other conditions were included: Condition 2 (Figure 2B) involved high-frequency word deviants embedded in a stream of well-matched pseudowords (HFW–PW), while condition 3 (Figure 2C) ivolved medium-frequency word deviants embedded in a stream of well-matched pseudowords (MFW–PW). Condition orders were counterbalanced across participants and classes. All three conditions were presented at a base frequency of 3 Hz and a deviant frequency of 1 Hz. This means that three stimuli per second were presented at a constant rate (3 Hz) and—for condition 1 for example—the three stimuli presented in a given second always comprised one learned low-frequency word (1 Hz) followed by two unlearned low-frequency words (LFW_UL_-LFW_UL_-LFW_L_). Of note, to avoid potential confounding of stimuli repetition in SSVEP studies (De Rosa et al., 2022), the pseudowords used in the HFW–PW condition were different from the pseudwords in the MFW–PW, which is why there are more pseudowords than word exemplars.

**Figure 2:**
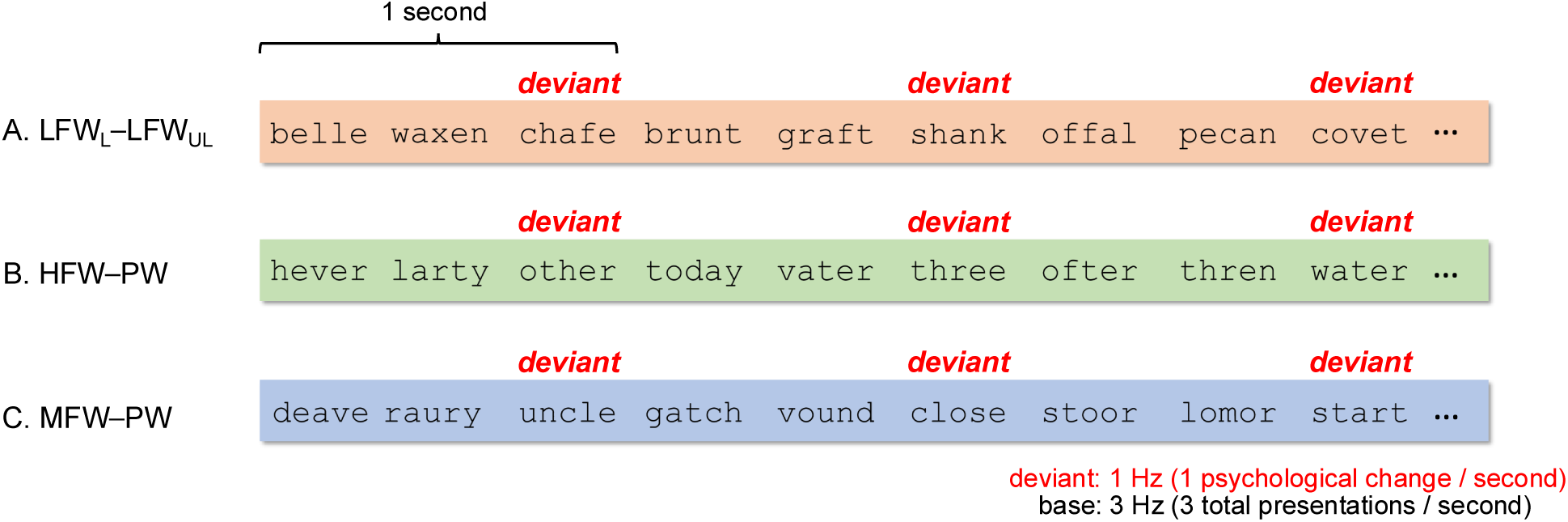
Experimental Design. Examples of stimuli presented in the experiment. 1 Hz deviants were embedded within a 3 Hz base stream in all three conditions. The first condition assessed processing differences between learned low-frequency words with well-matched unlearned low-frequency words (A. LFW_L_-LFW_UL_). The second condition assessed processing differences between high-frequency words with well-matched pseudowordss (B. HFW–PW). The third condition assessed processing differences between medium-frequency words with well-matched pseudowords (C. MFW–PW).

### 2.5 Behavioral Lexical Decision Task

All the stimuli in the lexical decision task were the same as those used in the EEG session. Children were asked to decide whether a stimulus was a real word or not. They pressed one button for a real word and one button for a pseudoword. The assignment to give the answer with the left or right index finger was balanced across subjects.

### 2.6 EEG Recording Procedure

Prior to the EEG recording, a brief practice session was held to familiarize the participant with the experimental procedure and the repetition detection task. During the EEG recording, participants sat in a dimly lit room 1 m away from the computer monitor. Each stimulation sequence started with a blank screen, the duration of which was jittered between 1500 ms and 2500 ms.

Each trial comprised 12 seconds of stimulation, and each condition comprised 10 trials (see Supplement Figure S1 for more information); thus, each participant completed 30 trials total across the three conditions. Participants were asked to press a button on an external response pad with their preferred hand when a stimulus (i.e., target) was repeated three times in a row.

Overall, the entire EEG experiment took around 40 minutes per participant including setup, practice, and breaks between trials and conditions.

### 2.7 EEG Recording and Preprocessing

EEG data were collected using 128-sensor HydroCell arrays (MagstimEGI), Electrical Geodesics NetAmp300, and NetStation 5.4.2 software, while stimuli were presented via an in-house software. Data were acquired against Cz reference at a sampling rate of 500 Hz. Impedances were kept below 50 kΩ.

Recordings were bandpass filtered offline (0.3–50 Hz) using Net Station Waveform Tools. Subsequently, data was preprocessed and re-sampled to 420 Hz. Sensors for which more than 15% of samples from the sensor exceeded a 60 *µ*V amplitude threshold were interpolated by the average value from six nearest neighboring sensors.

The continuous EEG data were then filtered with Recursive Least Squares (RLS) filters (Tang & Norcia, 1995) and re-referenced to average reference (Lehmann & Skrandies, 1980). Segented into 1-second epochs, data were screened for artifacts, excluding epochs with over 10% of data samples exceeding the noise threshold of 30 *µ*V or any part surpassing the 60 *µ*V blink threshold on a sensor-by-sensor basis. If an epoch exceeded the peak/blink threshold in more than 7 sensors, the entire epoch would be removed in all sensors. To mitigate initial transient responses, the first and last epochs of each 12-epoch, 12-second trial were omitted, leaving 10 epochs (i.e., 10 seconds) per trial for analysis.

The RLS filters were tuned to each of the analysis frequencies (base harmonics: 3Hz, 6Hz, 9Hz; deviant harmonics, excluding base harmonics: 1Hz, 2Hz, 4Hz, 5Hz, 7Hz, 8Hz) and converted to the frequency domain by means of Fourier transform. Complex-valued Fourier coefficients were decomposed into real and imaginary coefficients for input to the spatial filtering computations of Reliable Components Analysis (RCA), as described below.

### 2.8 Analysis of EEG Data

#### 2.8.1 Reliable Components Analysis (RCA)

We applied Reliable Components Analysis (RCA; Dmochowski et al. (2012, 2015)) to decompose the 128-sensor array into a set of *reliable components* (RCs) maximizing between-trials covariance. SSVEP response phases remain constant across stimulations, rendering RC activations indicative of phase-locked activities. RCA operates on sensor-by-feature EEG data matrices, deriving linear spatial filters to maximize Pearson Product Moment Correlation Coefficients (Pearson, 1896) across trials. These filters transform data from sensor-by-feature to component-by-feature matrices, with each component representing data from a linear combination of sensors. RC scalp topographies are visualized using forward-model projections of spatial filter vectors (Parra et al., 2005). Additional details on this spatial filtering technique are are provided in Dmochowski et al. (2012, 2015).

#### 2.8.2 RCA Calculations

To test whether low-level stimulus features were well matched across conditions, we conducted RCA at the base frequency and its harmonics. For this, we input to RCA the real and imaginary frequency coefficients for the first three harmonics (3 Hz, 6 Hz, and 9 Hz) across the 128-sensor array.

We next performed RCA at the deviant frequency and its six harmonics to examine processing disparities between deviants and control stimuli (i.e., LFW_L_ vs. LFW_UL_, HFW vs. PW, and MFW vs. PW). This encompassed real and imaginary Fourier coefficients at six harmonics, excluding base harmonics (1 Hz, 2 Hz, 4 Hz, 5 Hz, 7 Hz, and 8 Hz). RCA weights were computed separately for each condition to reveal potential similarities or differences underlying different stimulus contrasts.

#### 2.8.3 Analysis of RCA Data

We first assessed the significance of each component’s eigenvalue coefficient using permutation tests. This involved creating null distributions by generating 1000 surrogate data records with randomized phase spectra for each trial in sensor space before RCA computation. For more information, see Wang et al. (2022). Subsequently, we determined the significance of each harmonic within significant components using Hotelling’s two-sample t^2^ tests (Victor & Mast, 1991). Data were projected through spatial filter vectors, then averaged across epochs and participants before statistical analysis. False Discovery Rate (FDR, Benjamini & Yekutieli (2001)) correction was applied for multiple comparisons. For base analyses, corrections were made for 27 comparisons (3 harmonics *×* 3 components *×* 3 conditions). For deviant analyses, corrections were applied for 6 comparisons (6 harmonics *×* 1 component) per condition. Statistically significant RCs, verified by permutation testing and containing significant amplitudes in at least one harmonic, were further analyzed and reported in the results.

#### 2.8.4 Visualization of RCA Data

For both the deviant and base analyses, we visualized the data in two ways. First, we present topographic maps for spatial filtering components. Second, we present bar plots of amplitudes (*µ*V) across harmonics, with significant responses (according to adjusted *p_F DR_* values of Hotelling’s t^2^ tests of the complex data) indicated with asterisks.

### 2.9 Assessing Brain-Reading Relationships

Brain-behavior analyses were performed to assess individual variations in the relationship between EEG amplitude with reading scores. The reading scores analyzed were the participants’ grade-scaled scores of TOWRE, RANcolor, RANletter, and WJ.

Here, we focused on brain responses to deviants, which indicate discrimination responses between deviants and control stimuli. For HFW–PW and LFW_L_– LFW_UL_ conditions, where the first and fourth harmonics (1 Hz and 4 Hz) were significant, linear correlations were performed on the averaged projected amplitude between these two significant harmonics. For MFW–PW we performed linear correlations on projected amplitudes only at 1 Hz, which was the only significant harmonic.

If significant correlation was found, influential data points were identified using Cook’s Distance (Cook, 1977) and removed if they exceeded the CooksD 4*/n* threshold (where *n* is the total number of data points). Correlation analyses were then performed again to determine whether the significant relationship still held after the removal of influential data points.

### 2.10 Analysis of Repetition Detection During EEG

For behavioral responses to the repetition detection task during EEG recording, we computed *d’* based on the z-transformed probabilities of hits and false alarms (Macmillan & Creelman, 2004). One-way ANOVA with within-factor of condition was computed on *d’* across three conditions.

## 3 Results

### 3.1 New word learning at behavioral level

For the lexical detection task, means and standard deviations, *m*(*SD*), of mean accuracy in each class for each word group (one group a given student learned and two groups they did not learn) are summarized in Figure 3.

**Figure 3:**
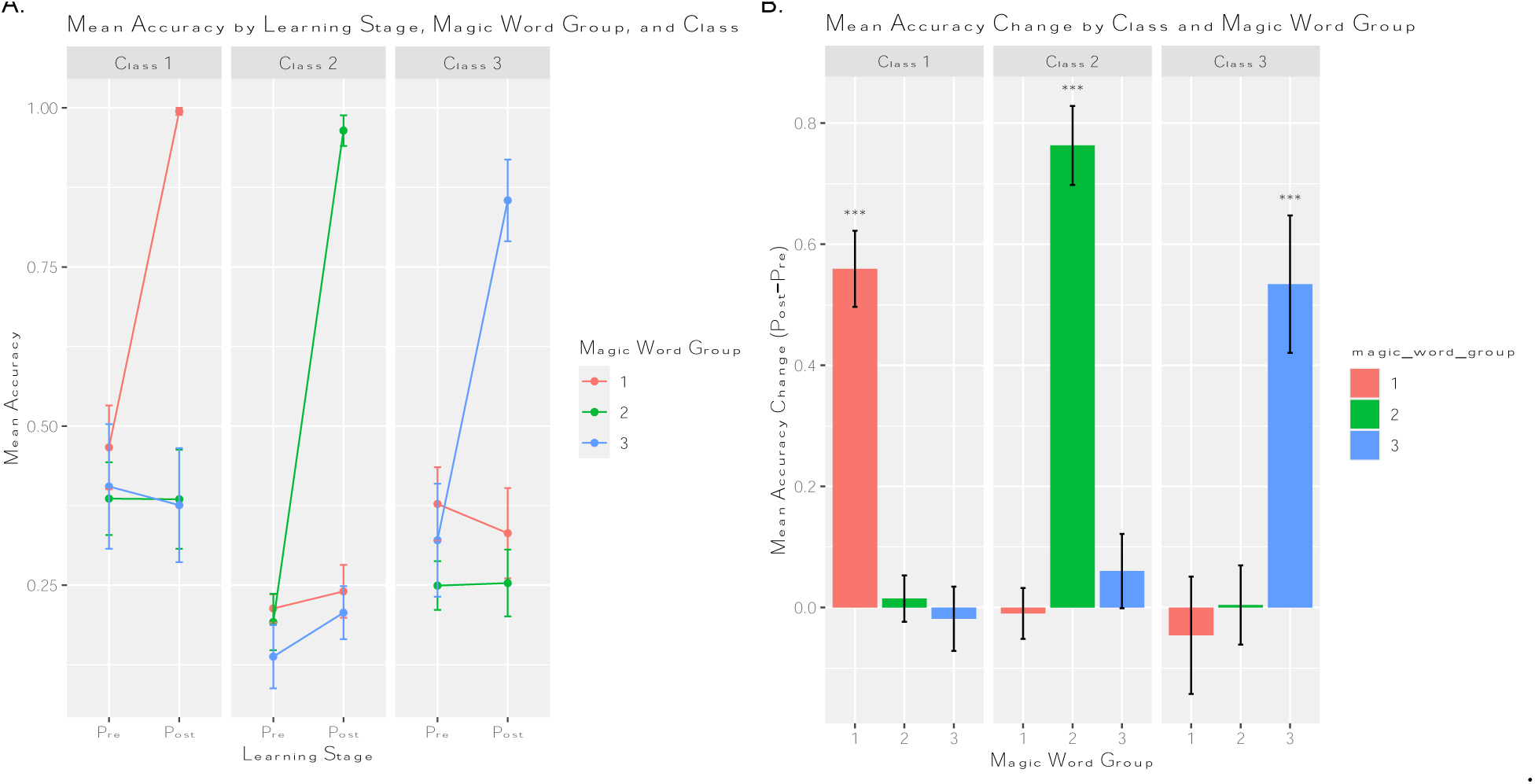
New word learning at behavioral level: Lexical decision task performance. A: Mean accuracy of behavioral responses to words that children learned in their own class and those they did not learn in the other two classes, before and after the two-week learning sprint; B: Mean accuracy changes (Post-Pre learning) of behavioral responses to words that children learned in their own class and those they didn’t learn in the other two classes. Children’s accuracy changed significantly for words they learned but not for those they didn’t learn.

A one-way ANOVA showed that mean accuracy did not differ significantly across the three conditions (*F* (2, 83) = 0.86, *p* = 0.43). Thus, we conclude that participants were equally engaged throughout the three conditions of the experiment. More importantly, for all three classes, mean accuracy changed significantly (all *p* < 0.001) for words students learned in their own class, but not for unlearned words.

### 3.2 Similar brain responses to high-frequency words and new learned words

Given the results from base frequencies reflect generalized visual processing, while the deviant results focus on the specific research questions you seek to address with each stimulus condition. Therefore, we will only present deviant analyses results in the main text, while keeping base analyses results in the supplement.

For responses to all three stimulus contrasts, only the first reliable component contained significant stimulus-driven activity (i.e., significant permutation test p-values and at least one significant harmonic). Results are summarized in Figure 4.

**Figure 4:**
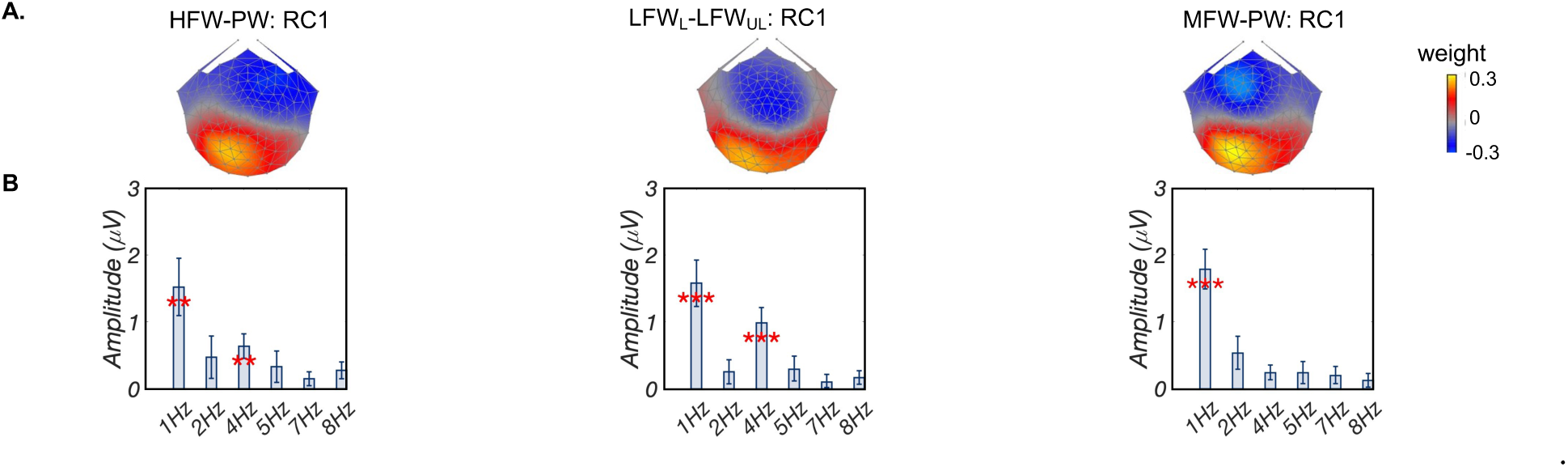
Comparison of brain responses to HFW–PW, LFW_L_– LFW_UL_, and MFW–PW. A: Topographic visualizations of the spatial filters for the first reliable component (RC1) in three conditions; B: Projected amplitude for each harmonic in bar charts. Only neural responses at 1 Hz and 4 Hz were significant. Responses at 2, 5, 7, and 8 Hz were not significant in neither of the three conditions. At 1 Hz, comparable responses amplitudes were found across three conditions. At 4 Hz, HFW–PW and LFW_L_– LFW_UL_ evoked significantly higher (both *p_F DR_* < 0.01) amplitude than MFW–PW; no significant difference was found between HFW–PW and LFW_L_–LFW_UL_. **: *p_F DR_* < 0.01, ***: *p_F DR_* < 0.001. HFW–PW: High-frequency words–Pseudowords; LFW_L_– LFW_UL_: Learned low-frequency–Unlearned low-frequency; MFW–PW: Medium-frequency–Pseudowords.

Figure 4A displays topographic visualizations of the spatial filter for RC1 in three conditions. RC1 topographies were highly correlated (*r* > 0.81) across the three conditions and maximal over left occipito-temporal area. The bar plots in Figure 4B present amplitudes, statistically significant responses were observed at first and fourth harmonics (all *p_F DR_* < 0.01, corrected for 6 comparisons) for HFW–PW and LFW_L_– LFW_UL_; but only at the first harmonic (*p_F DR_* < 0.001, corrected for 6 comparisons) in MFW–PW.

One-way ANOVA of response amplitudes at the first harmonic (1 Hz) did not show a main effect of condition (*F* (2, 83) = 0.15, *p* = 0.86), indicating comparable response amplitudes at the first harmonic across three conditions. However, one-way ANOVA at the fourth harmonic (4 Hz) showed a significant condition effect (*F* (2, 83) = 4.24, *p* < 0.05). Post-hoc paired t tests (one tailed) showed that response amplitudes at 4 Hz in conditions HFW–PW and LFW_L_– LFW_UL_ were significantly larger (both *p_F DR_* < 0.01, corrected for three comparisons) than that in MFW–PW condition. No significant difference (*t*(27) = 1.27, *p* = 0.11) was found between HFW–PW and LFW_L_– LFW_UL_. These results indicated similar responses between HFW and LFW_L_.

### 3.3 Individual differences in new vocabulary learning

For contrast of LFW_L_–LFW_UL_, we found significant correlations between EEG amplitudes at 1 Hz and both word decoding ability (measured by Woodcock-Johnson, WJ, *R*^2^ = 0.27, *p* < 0.01) and rapid naming ability for color (RANcolor, *R*^2^ = 0.3, *p_F DR_* < 0.01). No significant correlations were found for amplitudes at 4 Hz (all *R*^2^ < 0.1, *p* > 0.11) in this contrast. For contrasts of HFW–PW and MFW–PW, no significant correlations between response amplitudes and reading scores were found (all *R*^2^ < 0.12, *p* > 0.1).

**Figure 5:**
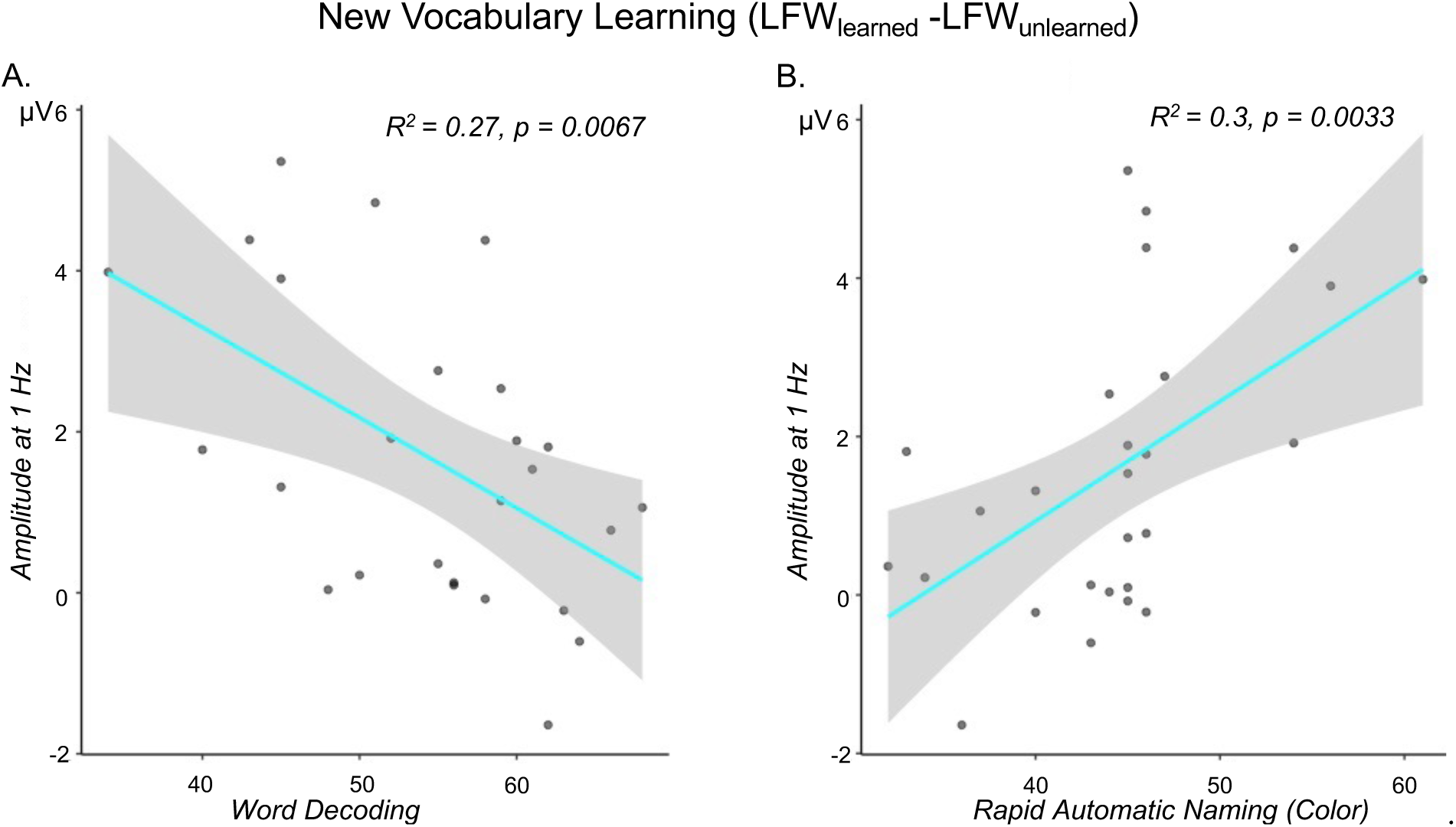
Statistically significant correlations between reading and new vocabulary learning effect, after outlier removal. A: Correlation between amplitudes at 1 Hz for LFW_L_–LFW_UL_ and word decoding ability (Woodcock-Johnson, WJ); B: Correlation between amplitude at 1 Hz for LFW_L_– LFW_UL_ and Rapid Automatic Naming (RAN, color). No significant relations were found for the other two contrasts: HFW–PW and MFW–PW. HFW: high-frequency words; MFW: medium-frequency words; LFW: low-frequency words; PW: pesudowords.

## 4 Discussion

In this study, we examined the question of whether neurocognitive responses to newly leaned words are similar to those elicited by words individuals have learned over their reading lifespan. We found that after two weeks of in-class learning, new vocabulary words evoked neural responses similar to high-frequency words, more so than medium-frequency words, reflecting learning at a higher-order lexical level instead of mere familiarity due to exposure. In addition, children with better phonological decoding skills (i.e., ability to sound out words) had more pronounced learning effect for new words. This study expands current understanding of word acquisition by highlighting the causal relationship between learning and word representation formation. Moreover, the in-school EEG-SSVEP paradigm permits short data collection sessions, paving the way for widespread adoption of educational neuroscience research within school environments.

### 4.1 Short-term learning builds lexical representations of new vocabulary words

RCA of learned low-frequency words in the LFW_L_–LFW_UL_ contrast generated a component that is distributed over the left occipito-temporal (OT) electrodes. This finding is in agreement with previous studies, in which robust training effects over the left OT are consistently revealed for whole word or single letter processing after short-term training. However, most previous training studies used stimuli from a novel language system (Brem et al., 2018; Hashimoto & Sakai, 2004) or an artificial script (Maurer et al., 2010; McCandliss et al., 1997; Pleisch et al., 2019; Xue et al., 2006; Yoncheva et al., 2010) in controlled lab settings, which limits the generalization to actual pedagogical practice and real vocabulary learning. For instance, with artificial grapheme-phoneme correspondence training, Pleisch et al. (2019) found more pronounced activation for trained than untrained *false-font* characters in the left OT. The current study extends previous work by demonstrating a learning effect for real 5-letter words—and crucially—in a natural school context instead of laboratory-based training experiments conducted previously. This contributes richly in bridging education professionals and researchers, as well as providing insights for developing effective learning curricula.

The present study additionally extends previous studies by comparing discriminative responses between learned real words vs. unlearned *real words* rather than *symbols* or *pseudowords*. The learned and unlearned words are well-matched in unigram, bigram, and trigram frequencies, as well as orthographic neighborhood sizes (see detailed statistics in Section 2.4 Stimuli). Such contrasts point towards a learning effect at the lexical level, instead of mere visual familiarity or mixed effects of learning at several levels of analysis (e.g., letter forms, sublexical and lexical information) that were captured in prior learned words vs. symbols contrasts.

### 4.2 Relationship between reading ability and new vocabulary word learning

We found that brain responses to learned words (vs. unlearned words) correlated with reading ability. Children with smaller response amplitudes distinguishing learned from unlearned words showed better word decoding and rapid automatic naming (RAN) skills, aligning with the importance of phonological decoding in learning novel words (Ziegler et al., 2014).

Word decoding can refer to the mapping of letters and groups of letters within words to their corresponding sounds—a process that encompasses a number of lexical and sublexical components including phonological awareness and orthographic processing (Ehri, 2005; Haft et al., 2019). For typical readers, word decoding has been used as a self-teaching strategy that establishes new written words in memory (Share & Shalev, 2004). However, many children with reading difficulties such as dyslexia have problems with word decoding, which in turn impacts the development of reading fluency (Nation & Snowling, 1997; Snowling, 2013). McCandliss and colleagues developed a decoding intervention program which focused on children’s attention to decoding each of the constituent grapheme–phoneme elements within words. They found that the decoding intervention had a substantial impact on reading-related skills—beyond decoding ability—in children with deficient decoding (McCandliss et al., 2003). Our finding, along with previous studies, supports the idea that word decoding ability plays a crucial role in word recognition (Al Otaiba et al., 2012) and reading acquisition (Stanovich et al., 1984).

The correlation between new vocabulary learning and reading ability was reinforced by its association with RAN. A task involving retrieval of associations between visual symbols and phonological codes (names), RAN predicts reading skills (Bowers & Newby-Clark, 2002; Georgiou et al., 2009; Schatschneider et al., 2004), and impacts literacy acquisition in elementary school children across different languages (Landerl et al., 2019; Moll et al., 2014). Some researchers consider RAN as a microcircuit of the later-developing reading circuitry (Lervåg & Hulme, 2009; Norton & Wolf, 2012). For instance, Lervåg & Hulme (2009) suggested that RAN taps object-naming circuits in the left hemisphere that are recruited to form the basis of the child’s developing visual word recognition system (Lervåg & Hulme, 2009).

Overall, correlation results in the current study provide further support for the idea that children rely on phonological decoding skills (the ability to sound out words/objects) to learn novel words.

### 4.3 Lexical cortical tuning for print in early readers

In addition to revealing fresh insights into vocabulary learning in natural classroom settings, we found that both high- and medium-frequency words, in contrast with pseudowords, evoked activation over left occipito-temporal regions, which has been challenging in previous SSVEP studies (Lochy et al., 2016; van de Walle de Ghelcke et al., 2021). We used a relatively explicit repetition detection task and slower presentation rates, potentially explaining this disparity. The task in our study may have directed more attention to linguistic aspects, unlike previous studies that may have focused more on implicit visual discrimination processes (e.g., color detection task in van de Walle de Ghelcke et al. (2021)). Additionally, the presentation rates used in previous studies (e.g., 1.2 Hz deviants and 6 Hz base in Lochy et al. (2016)) might have been too high for early readers, impacting higher-order processes. Our findings of lexical cortical tuning to print over left OT in early readers might reflect either direct access to lexical representations as indicated by numerous fMRI studies showing that left vOT regions, especially anterior, are engaged in lexical processing (Brem et al., 2010; Lerma-Usabiaga et al., 2018; Vinckier et al., 2007). Alternatively, it may also reflect indirect lexical access through phonological decoding or grapheme-phoneme mapping, as suggested by phonological mapping hypothesis (Maurer & McCandliss, 2007). As an example, a combined EEG/fMRI study showed that activation of VWFA is responsible for gapheme-to-phoneme conversion, especially during early reading phases when direct lexical mapping of orthographic information has yet to be built (Brem et al., 2010).

### 4.4 An SSVEP word frequency effect

A large number of studies have examined word frequency effects across different writing systems using different neuroimaging techniques (Assadollahi & Pulvermüller, 2003; Brysbaert et al., 2018; Grainger et al., 2012; Hauk & Pulvermüller, 2004; Strijkers et al., 2015; Zhang & Wang, 2014). Low-frequency words consistently elicit larger EEG/MEG responses and stronger fMRI activations over the left vOT compared to high-frequency words, aligning with behavioral findings showing shorter reading time for high-frequency words (Gregorová et al., 2023; Sereno & Rayner, 2003).

The current study, for the first time, examined the word frequency effect using an SSVEP paradigm. We contrasted high- and medium-frequency words separately with pseudowords to examine lexical retrieval. In comparison with a medium-frequency words vs. pseudowords contrast, the high-frequency words vs. pseudowords contrast elicited larger amplitude responses, particularly at the fourth harmonic (i.e., 4 Hz), suggesting facilitated access to lexical representations compared to medium-frequency words (Degani et al., 2019; McDonald & Shillcock, 2001; Ranbom & Connine, 2007). However, our study lacked a direct comparison between high- and medium-frequency words. Future studies with this stimulus contrast could further elucidate present findings.

### 4.5 Classroom Training Study Through a Research-Practice Partnership

Our research team, in partnership with a local elementary school, bridges the gap between education and neuroscience through collaborative studies in real-world educational settings. This enduring partnership facilitates classroom-based training studies, where educators co-design teaching activities and strategies. Additionally, the partnership includes a permanent on-site EEG recording studio at the school, enabling direct monitoring of how schooling impacts children’s brain development. Despite challenges, this collaboration enhances the practical application of science in schools, benefiting teachers, students, and researchers alike.

## 5 Conclusion

In summary, the present study revealed causal relationship between short-term classroom-based learning and word lexical representation building via a research-practice partnership between a university research team and a local school. This was demonstrated by the emergence, after two weeks of classroom-based learning, of discriminative responses between learned low-frequency words and unlearned low-frequency words, which are similar to the response pattern in the contrast of high-frequency words vs. well controlled pseudowords. In addition, we also found significant correlations between new vocabulary learning and reading skills, including word decoding and rapid automatic naming. The correlation results provided support for the notion that children rely on phonological decoding skills to learn novel words. Furthermore, responses to high-frequency words were higher than that to medium-frequency words, replicating the classic word frequency effect. Taken together, the present results from a natural school context extend previous laboratory-based knowledge on learning of new vocabulary words. Research in this direction could be utilized to develop effective learning curricula. The RPP model can be employed to conduct neuroscience in ecologically valid educational settings, drawing more direct connections between education and neuroscience.

## Conflict of Interest Statement

The authors declare that the research was conducted in the absence of any commercial or financial relationships that could be construed as a potential conflict of interest.

## Data Availability Statement

The data sets generated for this study are available upon reasonable request to the corresponding author.

## Funding Statement

This research did not receive any specific grant from funding agencies in the public, commercial, or not-for-profit sectors

## Author Contributions

F.W. and B.D.M. conceived the study. F.W., E.Y.T., and R.S.G conducted the experiment(s). F.W. analyzed the data and wrote the original draft. B.K., A.M.N. B.D.M. edited the draft. All authors reviewed the manuscript.

## Ethics Approval Statement

The study was approved by the Institutional Review Board of Stanford University. A parent or legal guardian of each participant received a written description of the study and gave written informed consent before the session; each participant also assented to participating.

## Acknowledgments

We thank the students, their families, and teachers for participating. We also thank Lindsey Hasak and Suanna Moron for their help with data collection.

## Footnotes

1 First and second graders were in the same class at the school.

## 1 Supplementary Material

### 1.1 Nontarget, terminal, and catch trials

Among the 10 trials that were pseudorandomly presented for each condition, there were four “nontarget” trials which contained no repeated stimuli (Figure S1A); four “terminal” trials in which repeated stimuli appeared at the end (Figure S1B); and two “catch” trials with repeated stimuli randomly appearing elsewhere during the trial (Figure S1C). Each terminal and catch trial contained only one target. Participants were given verbal feedback about their performance after the end of each trial. Due to excessive movements from the button press, each participant’s two catch trials were excluded in their entirety from further analysis. Data corresponding to the four terminal trials were still included because movements happened at the end of the trial. In all, 8 of 10 trials for each condition from each participant were used for analysis.

**Figure S1:**
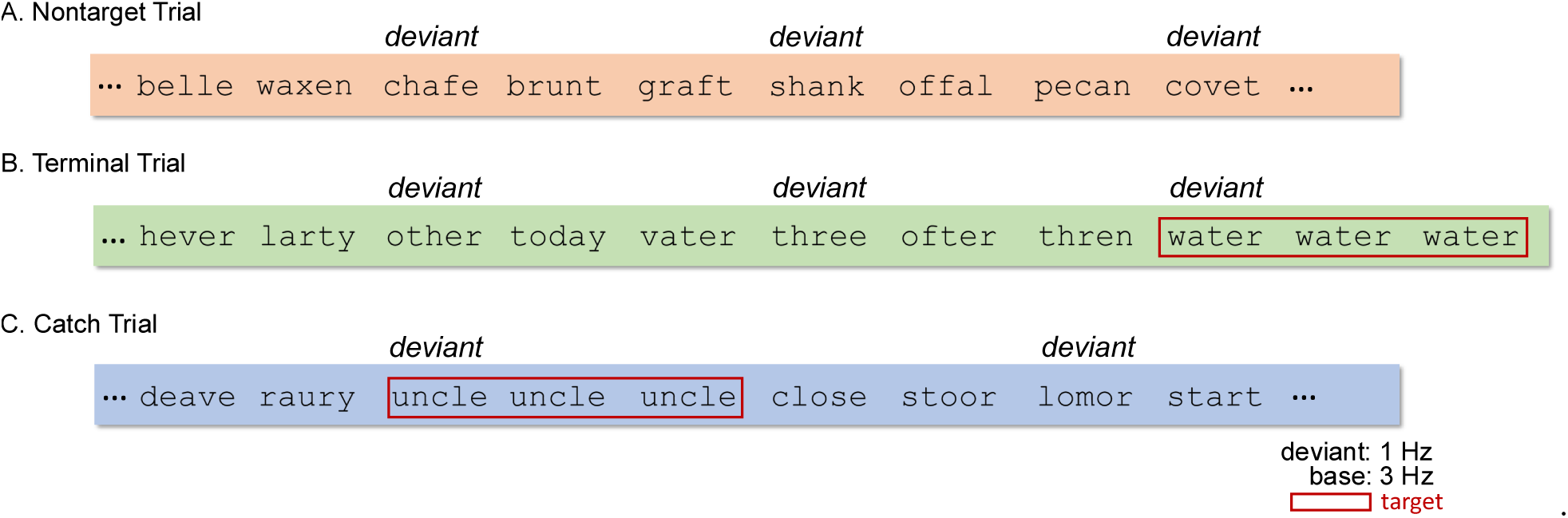
Example nontarget, terminal, and catch trials. Twelve trials were pseudorandomly presented for each condition, including four nontarget trials (A), four terminal trials (B), and two catch trials (C). Responses to all catch trials were excluded from EEG analyses due to excessive response-related movements during recording. Data corresponding to the four terminal trials were still included because movements happened at the end of the trial, resulting in 8 out of 10 trials per condition and participant being used for analyses.

### 1.2 Base Analyses Results

We performed RCA on responses at the base frequency and its harmonics in order to investigate neural activity related to low-level visual processing and whether visual properties of different types of stimuli were well matched across conditions. Based on permutation test and Hotelling’s t^2^ test, we report spatial filter topographies and statistical analysis of the projected data for the first three RCs (RCs 1, 2, and 3) returned by RCA trained on three conditions together. Base RCA results are summarized in Figure S2.

Figure S2A displays topographic visualizations of the spatial filters for the first three reliable components that contained significant stimulus-driven activity (i.e., significant permutation test and at least one harmonic is significant). Figure S2B shows summary plots of the responses in the 2D complex plane, with overlapping amplitudes (vector lengths) and phases (vector angles) between these three conditions. Figure S2C presents projected amplitude (project sensor-space data through spatial filter) in bar plots and the derived latency estimations. The projected amplitude contained statistically significant responses in all three harmonics at each of the three RCs (all *p_F DR_* < 0.05, corrected for 27 comparisons) for all three conditions.

*RSS* amplitude (across three significant harmonics) comparisons showed that there is no significant difference between conditions at each of three RCs (all *F* (2, 83) < 0.35, all *p* > 0.70). Latency estimation derived from phase slopes across significant harmonics were also similar between conditions (RC1: 147.99 ms, 147.29 ms, and 148.29 ms; RC2: 309.68 ms, 313.33 ms, and 314.38 ms; RC3: 84.28 ms, 93.65 ms, and 92.98 ms; for three conditions respectively).

### 1.3 Participants were equally engaged throughout the experiment

For the repetition detection task, means and standard deviations, *m*(*s*), of *d’* were 1.04(0.38), 1.18(0.43), 1.14(0.40) for LFW_L_– LFW_UL_, HFW–PW, and MFW–PW, respectively. A one-way ANOVA showed that *d’* did not differ significantly across the three conditions (*F* (2, 83) = 0.86, *p* = 0.43). Thus, we conclude that participants were equally engaged throughout the three conditions of the experiment.

**Figure S2:**
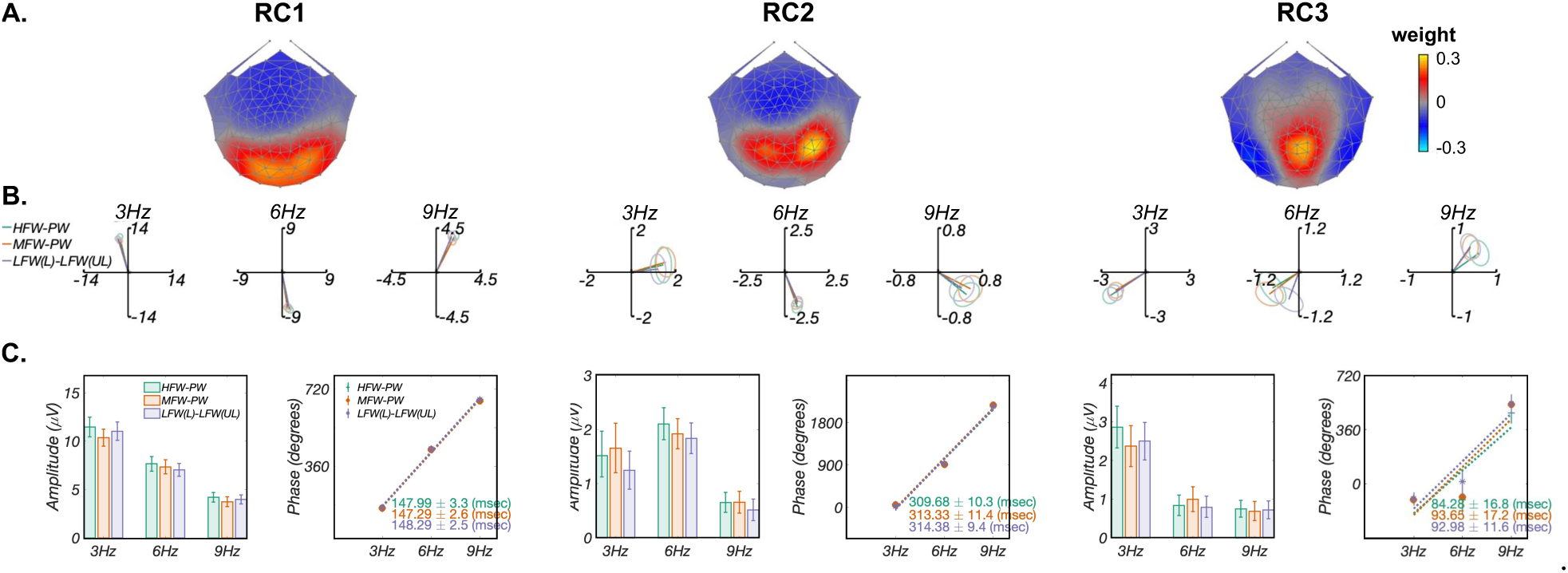
Base Analyses Results. A: Topographic visualizations of the spatial filters for the first three components (RC1, RC2, and RC3); B: Response data of three conditions presented in the 2D complex plane, where amplitude information is represented as the length of the vectors, and phase information in the angle of the vector relative to 0 degrees (counterclockwise from 3 o’clock direction), ellipse indicates standard error of the mean (*SEM*); C: Comparison of projected amplitude and latency estimation across three conditions. The projected amplitude contained statistically significant responses in all three harmonics at each of the three RCs (all *p_F DR_* < 0.05, corrected for 27 comparisons) for all three conditions. In addition, *RSS* response amplitudes across three harmonics did not differ significantly across three conditions for RCs 1-3 (all *F* (2, 83) < 0.35, all *p* > 0.70), latency estimations derived from phase slopes across harmonics were also similar across conditions.

## Notes

### Competing Interest Statement

The authors have declared no competing interest.

